# Ecological Toxicity Effects of Marine Pollutants Based on Microalgal Locomotory Behavior

**DOI:** 10.1101/2025.04.23.650213

**Authors:** Zhanfei Xie, Lin Wang, Guoxia Zheng

**Author notes:** Corresponding Author E-mail addresses.

## Abstract

Understanding the ecological toxicity of marine pollutants is essential for assessing marine environmental quality and developing effective protection strategies. This study presents a novel approach using the motility of the marine microalgae as a sensitive physiological indicator for rapid toxicity assessment. Results demonstrated that both copper (0µmol l^-1^-4.4µmol l^-1^) and phenol (0mmol l^-1^-9.1mmol l^-1^) significantly inhibited algal locomotory behavior parameters (movement patterns, mobility, and swimming speed) within 2 hours, exhibiting clear dose-response relationships. The 2 h-EC50 values determined by logistic regression were 2.2µmol l^-1^-2.7µmol l^-1^ for copper and 4.5µmol l^-1^-5.7 mmol l^-1^ for phenol. Notably, combined exposure showed antagonistic effects (2 h-EC50 >1 TU). The motility-based assay showed strong correlation with traditional toxicity tests (72h growth inhibition, 24-48 h *Daphnia* immobilization, and 96h fish mortality), while providing significant advantages in speed and sensitivity. These findings establish microalgal motility as an efficient, ecologically relevant biomarker for comprehensive marine pollutant assessment, offering great potential for environmental monitoring and risk evaluation.

## 1 Introduction

The accelerating degradation of marine ecosystems under increasing anthropogenic pressures has become a critical environmental concern worldwide. Among diverse marine contaminants, phenolic compounds and heavy metals represent two major classes of pollutants with distinct yet equally concerning ecological impacts. Phenol, as the simplest aromatic alcohol and a fundamental unit of more complex phenolic pollutants, exhibits widespread distribution in coastal environments [1]. Similarly, copper, while biologically essential as a micronutrient, demonstrates pronounced ecotoxicity at elevated concentrations typical of polluted marine systems [2]. These contaminants differ fundamentally in their chemical behavior, bioavailability, and mechanisms of toxicity, yet frequently co-occur in marine environments, creating complex interactive effects that challenge conventional risk assessment paradigms [3].

Primary producers, particularly marine microalgae, occupy a pivotal position in marine ecosystem vulnerability to pollutants. As the foundation of pelagic food webs and key contributors to global carbon cycling, microalgae serve as sensitive bioindicators of marine environmental health. Traditional ecotoxicological assessments have relied on physiological endpoints including growth inhibition, photosynthetic efficiency, and biochemical markers [4, 5]. However, these approaches often require extended exposure periods and may overlook subtle yet ecologically significant sublethal effects.

The locomotory behavior of marine phytoplankton presents an underutilized but highly informative biomarker that bridges cellular physiology with ecosystem-level consequences. Flagellar motility, exhibited by over 90% of harmful algal bloom species [6-7], mediates critical ecological processes including vertical migration, niche partitioning, and predator-prey interactions [8-11]. As a distinctive physiological characteristic, microalgal locomotory behavior shares similarities with other behavioral indicators used in ecotoxicological assessments, such as fish gill movements, avoidance behaviors, Daphnia swimming patterns, and feeding behaviors. These behavioral endpoints offer intuitive and observable responses to environmental stressors which hold unique advantages as an ecotoxicological endpoint: (1) rapid response times reflecting immediate physiological stress, (2) high sensitivity to sublethal contaminant levels, and (3) direct ecological relevance to population dynamics and community structure.

Despite these advantages, the application of microalgal locomotory behavior in ecotoxicological assessment remains surprisingly limited. We address this knowledge gap through a comprehensive investigation using the model species *Platymonas subcordiformis*, an ecologically relevant, motile microalga prevalent in Chinese coastal waters. Our study systematically evaluates: (1) concentration-response relationships for individual copper and phenol exposure, (2) interactive effects in combined exposure scenarios, and (3) the ecological implications of contaminant-induced motility alterations. The results establish microalgal locomotory behavior as a robust, rapid, and ecologically grounded biomarker for marine pollution assessment, with particular relevance for developing early warning systems in vulnerable coastal ecosystems.

## 2 Materials and methods

### 2.1 Test algal strain and culture

The marine microalga *Platymonas subcordiformis* was obtained from the Institute of Oceanology, Chinese Academy of Sciences. Cultures were maintained in f/2 medium at 20±0.5°C under 55 μmol (m^2^·s)^-1^ illumination with a 12h : 12h light-dark photoperiod. Algal cells in the early exponential growth phase were selected for toxicity experiments.

### 2.2 Single toxicity tests of copper and phenol

Preliminary experiments were conducted to optimize concentration ranges and minimize errors in calculating 2h EC_50_ values. Based on these results, test concentrations for copper and phenol were established at arithmetic intervals: Copper: 0 (control), 0.6, 1.3, 1.9, 2.5, 3.2, 3.8, and 4.4µmol l^-1^; Phenol: 0 (control), 1.3, 2.6, 3.9, 5.2, 6.5, 7.7, and 9.1mmol l^-1^. All treatments were performed in triplicate using 96-well plates with 200 μl working volume at an initial cell density of 1×10^6^ cells ml^-1^. After 2h exposure, 30 μl aliquots were transferred to microscope slides, covered with coverslips, and observed under a Nikon 5100 microscope (100× magnification). Random fields of view were recorded for 30 s using a CCD camera for subsequent motility analysis.

### 2.3 Combined Toxicity Tests of Copper and Phenol

The combined toxicity experiments followed the method described by Charles *et. al*. [12]. Based on the EC_50_ values obtained from single toxicity tests (Section 2.2), one toxicity unit (TU) was defined as: 0.5×EC_50_ (phenol) + 0.5×EC_50_ (copper) = 1 TU. The combined toxicity test concentrations were set as: 0 (control), 0.3, 0.7, 1.0, 1.3, 1.6, 1.9, and 2.3 TU. All treatments were performed in triplicate using identical experimental conditions as described in Section 1.2 (96-well plates, 200 μl volume, 1×10^6^ cells ml^-1^). After 2h exposure, 30 μl samples were prepared for microscopic observation and video recording (Nikon 5100, 100× magnification) to analyze algal Locomotory parameters.

### 1.4 Data analysis and processing

The 30-second video recordings collected from toxicity tests (Sections 2.2 and 2.3) were analyzed using ImageJ software with the CASA (Computer-Assisted Sperm Analysis) plugin to calculate microalgal Locomotory parameters, including: Motility percentage (MOT%): Percentage of motile cells relative to total cells; Curvilinear velocity (VCL, μm s^-1^): Average velocity along the actual swimming path; Average path velocity (VAP, μm s^-1^): Velocity calculated from a smoothed movement trajectory; Straight-line velocity (VSL, μm s^-1^): Velocity measured from the straight-line distance between start and end points. The Locomotory parameters were plotted against pollutant concentrations and fitted using a Logistic model to quantify the response of *P*. *subcordiformis* to toxicants: f(x) = a / [1 + e ^(b(x-x0))^]

Where f(x) is locomotory behavior parameter (MOT%, VCL, VAP, or VSL) in treated groups; x is pollutant concentration; a is locomotory behavior parameter in control groups; b is slope parameter of the curve; x0 is pollutant concentration when motility decreases to 50% of control (EC_50_ value); The limit of detection (LOD) was defined as the pollutant concentration corresponding to a 10% reduction in motility compared to controls. Combined toxicity effects were evaluated using the Additive Index method ^[12]^, where: EC_50_ < 1 TU indicates synergistic effects; EC_50_ > 1 TU indicates antagonistic effects; EC_50_ = 1 TU indicates simple additive effects.

All experimental results were expressed as mean ± standard deviation (n=3). Statistical significance between treatment groups and controls was determined using Student’s t-test, with P<0.05 considered statistically significant.

## 3 Results and discussion

### 3.1 Response of microalgal locomotory patterns to pollutants

The effects of pollutants on the locomotory patterns of *P. subcordiformis* are illustrated in figure 1. Each black trajectory in the figure represents the swimming path of an individual microalgal cell. Numerous studies have confirmed that phytoplankton within the marine euphotic zone are predominantly motile ^[6,7]^. These microorganisms propel themselves by beating one or multiple flagella, exhibiting either linear or tumbling movements, which collectively manifest as random, non-directional swimming behavior ^[13]^. As shown in the control group trajectories (Fig. 1), *P. subcordiformis* displayed these characteristic movement patterns under pollutant-free conditions.

**Fig. 1.**
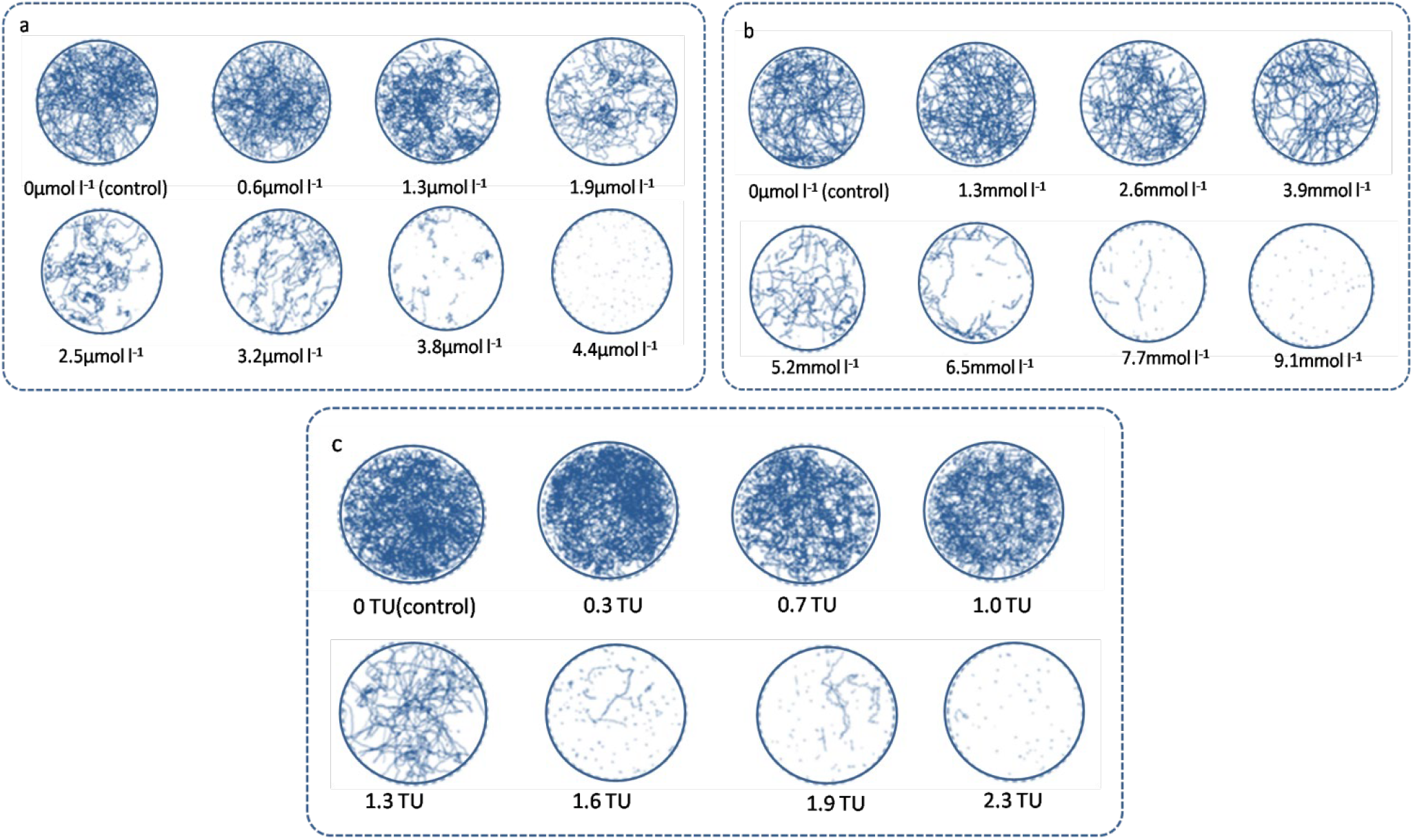
The effects of pollutants (a) Cu, (b) phenol and (c) their combines on the locomotory patterns of *P. subcordiformis*.

However, when exposed to copper, phenol, or their combination, the algae exhibited significant locomotory impairments characterized by progressively deteriorating movement coordination and swimming performance. Initial exposure resulted in disrupted movement patterns, with normally smooth tumbling motions becoming increasingly erratic. As pollutant concentrations increased, a clear dose-dependent reduction in swimming velocity emerged, evidenced by decreasing displacement per unit time. The most severe toxic effects manifested at higher concentrations, where cells displayed markedly constrained movement patterns ranging from unproductive circular swimming and abnormal trembling to complete immobilization. These behavioral modifications strongly suggest direct interference with the flagellar propulsion system by both contaminants, with the progressive nature of motility inhibition following classical toxicological response patterns. Notably, the sensitivity of these movement alterations to sublethal contaminant levels positions them as valuable visual indicators of physiological stress, offering real-time insights into pollutant impacts on microalgal function. The distinct stages of behavioral deterioration - from subtle coordination deficits to complete motility cessation - provide a quantifiable continuum for assessing toxicity thresholds and mechanisms.

### 3.2 Individual toxicity of phenol and copper

The dose-response relationships of copper and phenol on *P. subcordiformis* motility were quantitatively analyzed using an integrated machine vision and computer-assisted cell tracking system. Locomotory parameters, including the percentage of motile cells (MOT%) and swimming velocity components (VCL, VAP, and VSL), were fitted against pollutant concentrations using a logistic regression model (figures 2 and 3). All locomotory parameters exhibited significant concentration-dependent responses within just 2 hours of exposure. Figures 2a and 3a demonstrate the inhibitory effects on MOT%, with complete motility cessation occurring at 3.8 μmol l^-1^ copper and 7.7 mmol l^-1^ phenol. Interestingly, copper exposure at 0.6 μmol l^-1^resulted in slightly enhanced swimming speeds (VCL, VAP, and VSL) compared to controls (P<0.05), suggesting a potential hormetic effect at subtoxic concentrations. However, this stimulatory effect reversed to significant inhibition at ≥1.3μmol l^-1^ copper. In contrast, phenol exposure showed consistently inhibitory effects across all tested concentrations, with statistically significant velocity reductions (P<0.05) observed at ≥2.6 mmol l^-1^. The 2-hour limit of detection (LOD) based on locomotory parameters ranged from 0.7-1.7μmol l^-1^ for copper and 1.4-3.0mmol l^-1^ for phenol, demonstrating the sensitivity of this behavioral assay for rapid toxicity assessment.

**Fig. 2.**
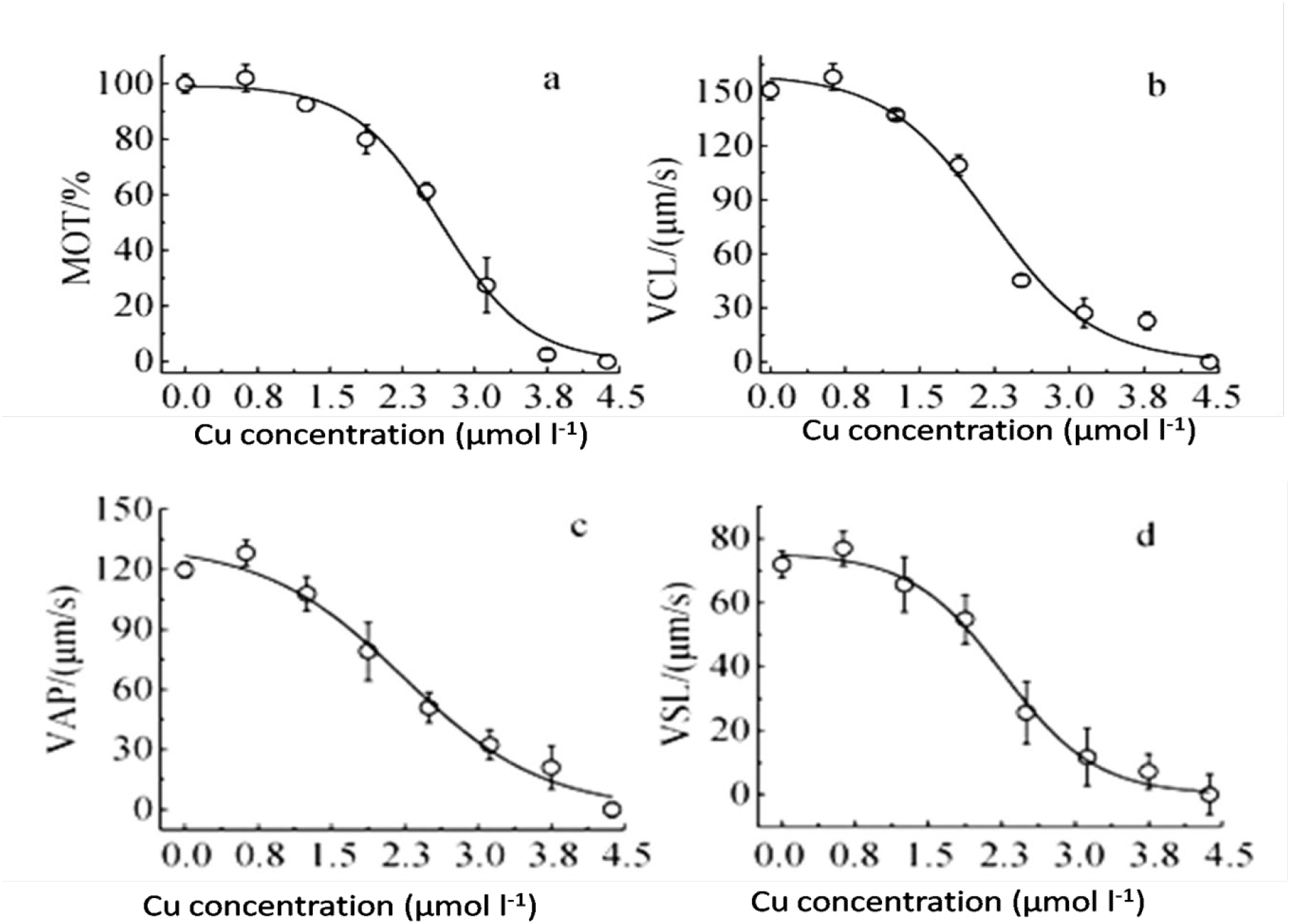
Locomotory parameters, including (a) the percentage of motile cells (MOT%) and swimming velocity components (b)VCL, (c)VAP and (d) VSL were fitted against Cu concentrations.

**Fig. 3.**
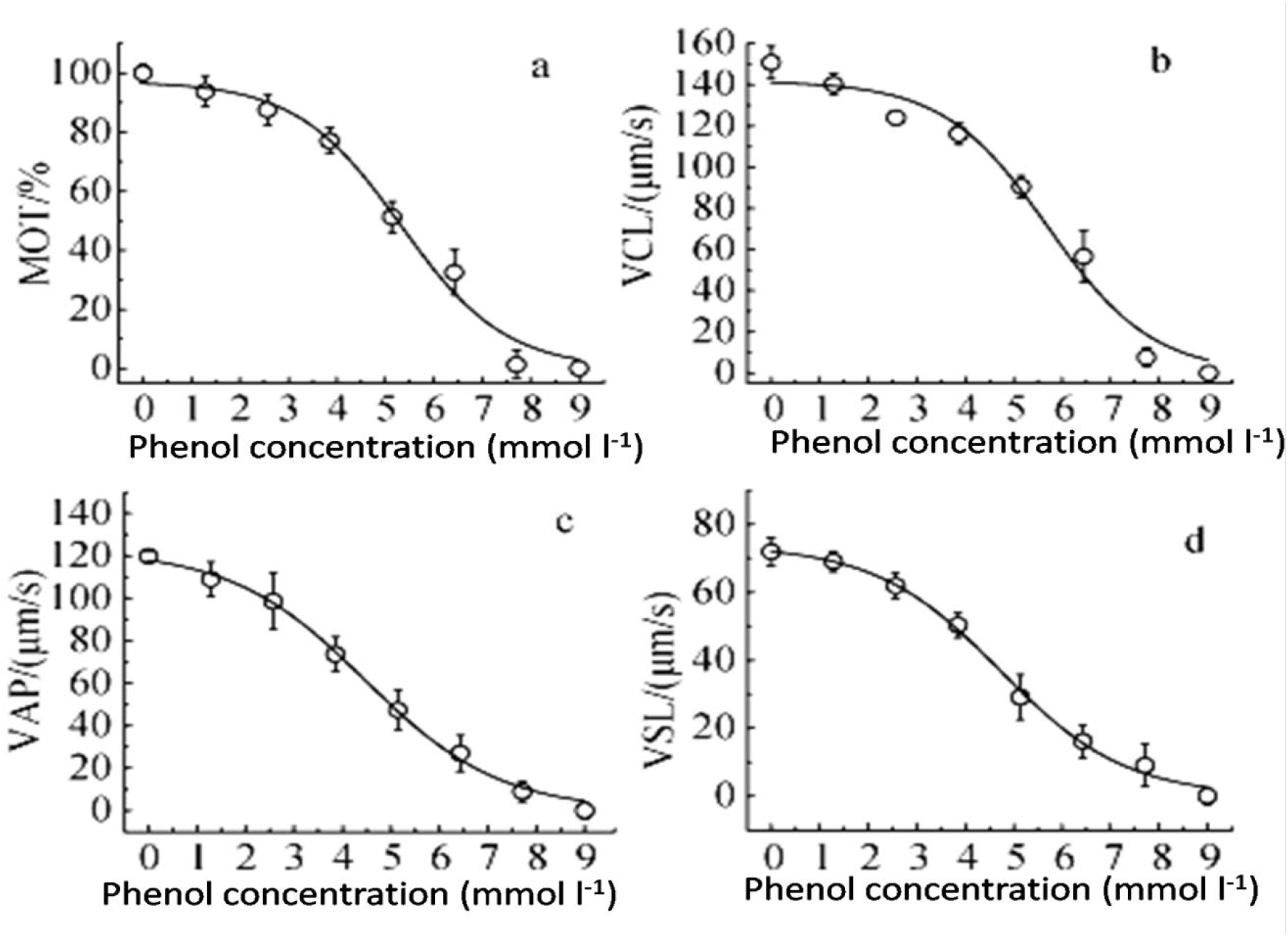
Locomotory parameters, including (a) the percentage of motile cells (MOT%) and swimming velocity components (b)VCL, (c)VAP and (d) VSL were fitted against phenol concentrations.

Flagella serve as the primary locomotory organelles in motile microalgae, propelling the cells through coordinated beating patterns of one to multiple flagella to achieve remarkable swimming speeds ranging from tens to hundreds of micrometers per second. In this study, *P. subcordiformis* exhibited an average swimming velocity of 151 μm s^-1^. Flagellar beating represents an energy-intensive process strictly dependent on ATP availability. Consequently, any factors disrupting ATP generation pathways whether through direct interference with photosynthetic electron transport chains, impairment of cellular catabolic processes, or indirect modulation of related signaling pathways, can significantly inhibit flagellar motility, rendering microalgal swimming behavior a sensitive toxicity indicator [14]. Different pollutants may share common or distinct molecular targets. Furthermore, certain contaminants may directly damage flagellar structures, leading to complete motility loss [15]. Specifically, Soretino et al. [16] demonstrated that copper ions can penetrate chloroplasts and uncouple photophosphorylation, thereby inhibiting photosynthesis. Pätsikkä et al. [17] further revealed that copper induces reactive oxygen species (ROS) generation, which damages the D1 protein in PSII oxygen-evolving complexes, ultimately disrupting electron transport and light reactions. Similarly, phenol predominantly exerts toxic effects through suppression of both photosynthetic and respiratory activities [18,19], with Zhang et al. [20] identifying particular vulnerability in dark reaction processes. For *P. subcordiformis*, both copper and phenol likely impair motility through these ATP synthesis disruption mechanisms.

The 2h EC_50_ values based on motility parameters were 2.2–2.7 μmol l^-1^ for copper and 4.5–5.7 mmol l^-1^ for phenol, showing strong consistency with conventional toxicity endpoints including 72 h algal growth inhibition (0.9-5.2μmol l^-1^), Daphnia immobilization (6.28μmol l^-1^), and fish mortality tests (4.72 μmol l^-1^) [21]-27]. Notably, the motility assay achieved comparable sensitivity within significantly shorter exposure periods. These findings establish microalgal locomotory parameters (including motility percentage and velocity components) as rapid, reliable biomarkers for marine pollutant assessment, offering substantial advantages for high-throughput environmental monitoring applications.

### 3.3 Combined toxic effects of phenol and copper

The combined toxic effects of phenol and copper on *P. subcordiformis* are presented in figure 4, showing the impact on motility percentage and swimming velocity parameters when mixed at an equitoxic ratio (1:1 TU). Logistic model fitting revealed 2 h EC_50_ values of 1.3-1.5 TU (Table 3), all exceeding 1 TU, indicating an antagonistic interaction between the two pollutants. When toxic substances are combined, the observed toxicity may differ from the expected additive effect through two primary mechanisms: first, chemical interactions between substances may alter their structures or properties, consequently modifying their biological toxicity; second, the mixture may influence the organism’s absorption, excretion or biotransformation of individual components. In the case of phenol-copper mixtures, the antagonism likely arises from the former mechanism. Typically, the toxicity of non-ionic organic compounds to aquatic organisms depends on their membrane permeability. Phenol can form phenol-copper-phenol complexes with copper ions, resulting in increased molecular size and reduced membrane penetration capacity [27]. Furthermore, the formation of these complexes decreases the concentration of biologically active free copper ions, leading to the observed antagonistic effect. This finding is consistent with results from 48 h Daphnia lethality tests [27], demonstrating the feasibility of using microalgal motility as a rapid screening tool for assessing combined toxicity of marine pollutants. The approach provides significant advantages in detection speed while maintaining ecological relevance, offering promising applications in marine environmental monitoring and risk assessment.

**Fig. 4.**
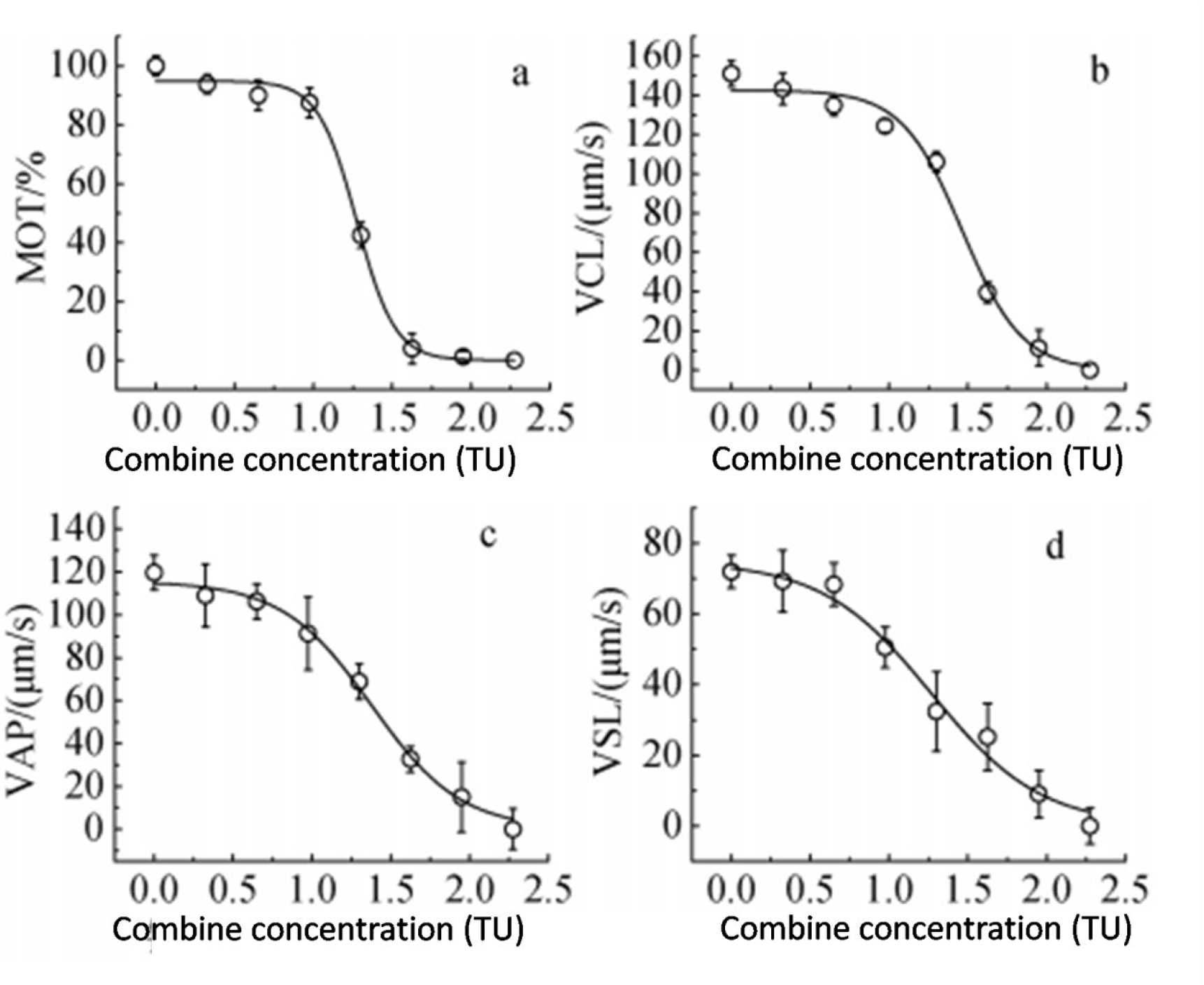
Locomotory parameters, including (a) the percentage of motile cells (MOT%) and swimming velocity components (b)VCL, (c)VAP and (d) VSL were fitted against phenol concentrations.

## 4 Conclusions

This study demonstrates that copper and phenol exposure, either individually or in combination, significantly impairs the locomotory behavior of Platymonas subcordiformis, as quantitatively characterized through integrated machine vision and computer-assisted cell tracking analysis. The microalgae exhibited dose-dependent motility inhibition within just 2 hours of exposure, progressing from reduced swimming speeds and disrupted movement patterns (including impaired rotation and circular swimming) to complete immobilization at higher concentrations. Quantitative assessment using multiple locomotory parameters (MOT%, VCL, VAP, and VSL) yielded 2 h-EC50 values of 2.2-2.7 μmol l^-1^ for copper and 4.5-5.7 mmol l^-1^ for phenol, while their combined exposure showed consistent antagonistic effects (EC50 >1 TU). The rapid response time, high sensitivity, and quantitative nature of this motility-based assay, coupled with its strong correlation with conventional toxicity endpoints, establish microalgal locomotion as a robust biomarker for marine pollutant assessment. These findings highlight the potential of this approach for both laboratory studies and field applications in environmental monitoring, particularly where rapid toxicity screening of multiple pollutant classes is required, though further validation across broader taxonomic groups and contaminant types would strengthen its generalizability as a standardized ecotoxicological tool.

## Declaration of competing interest

The authors declare that they have no known competing financial interests or personal relationships that could have appeared to influence the work reported in this paper.

## Acknowledgements

This work was supported by General program of Liaoning Science and Technology project (No. 2021-MS-345).

## Notes

### Competing Interest Statement

The authors have declared no competing interest.

